# Pneumococcal metabolic adaptation and colonization is regulated by the two-component regulatory system 08

**DOI:** 10.1101/300095

**Authors:** Alejandro Gómez-Mejia, Gustavo Gámez, Stephanie Hirschmann, Viktor Kluger, Hermann Rath, Sebastian Böhm, Franziska Voss, Niamatullah Kakar, Lothar Petruschka, Uwe Völker, Reinhold Brückner, Ulrike Mäder, Sven Hammerschmidt

**Affiliations:** Department of Molecular Genetics and Infection Biology, Interfaculty Institute for Genetics and Functional Genomics, Center for Functional Genomics of Microbes, University of Greifswald, Greifswald, Germany; Genetics, Regeneration and Cancer (GRC) Research Group, University Research Center (SIU), Universidad de Antioquia, UdeA, Medellin, Colombia; Basic and Applied Microbiology (MICROBA) Research Group, School of Microbiology, Universidad de Antioquia, UdeA, Medellin, Colombia; Department of Functional Genomics, Interfaculty Institute for Genetics and Functional Genomics, Center for Functional Genomics of Microbes, University Medicine Greifswald, Greifswald, Germany; Department of Microbiology, University of Kaiserslautern, Kaiserslautern, Germany

**Author notes:** Address correspondence to Prof. Dr. Sven Hammerschmidt.

## Abstract

*Streptococcus pneumoniae* two-component regulatory systems (TCS) enable adaptation and ensure its maintenance in host environments. This study deciphers the impact of the TCS08 on pneumococcal gene expression and its role in metabolic and pathophysiological processes. Transcriptome analysis and real-time PCR demonstrated a regulatory effect of the TCS08 on genes involved mainly in environmental information processing, intermediary metabolism, and colonization by *S. pneumoniae* D39 and TIGR4. Striking examples are genes of the fatty acid biosynthesis, arginine-deiminase system, and *psa* operon encoding the manganese ABC transport system. *In silico* analysis confirmed that TCS08 is homologous to *Staphylococcus aureus* SaeRS and a SaeR-like binding motif is displayed in the promotor region of *pavB*, the upstream gene of the *tcs08* operon encoding a surface-exposed adhesin. Indeed, PavB is regulated by the TCS08 as confirmed by immunoblotting and surface abundance assays. Similarly, Pilus-1 of TIGR4 is regulated by TCS08. Finally, *in vivo* infections using the acute pneumonia and sepsis models showed a strain dependent effect. Loss of function of HK08 or TCS08 attenuated D39 virulence in lung infections. The RR08 deficiency attenuated TIGR4 in pneumonia, while there was no effect on sepsis. In contrast, lack of HK08 procured a highly virulent TIGR4 phenotype in both pneumonia and sepsis infections. Taken together, these data indicate the importance of TCS08 in pneumococcal fitness to adapt to the milieu of the respiratory tract during colonization.

**IMPORTANCE:** *Streptococcus pneumoniae* interplays with its environment by using 13 two-component regulatory systems and one orphan response regulator. These systems are involved in the sensing of environmental signals thereby modulating pneumococcal pathophysiology. This study aimed to understand the functional role of genes subject to control by the TCS08. The identified genes play a role in transport of compounds such as sugars or amino acids. In addition, the intermediary metabolism and colonization factors are modulated by TCS08. Thus, TCS08 regulates genes involved in maintaining pneumococcal physiology, transport capacity and adhesive factors to enable optimal colonization, which represents a prerequisite for invasive pneumococcal disease.

## INTRODUCTION

Regulatory systems are inherent features of living organisms, ensuring a rapid response and adaptation to diverse environmental conditions and acting as on/off switches for gene expression (1). Regulation in bacteria is predominantly conducted by two-component regulatory systems (TCS), quorum sensing proteins and stand-alone regulators (2–4). TCS are the most common and widespread sensing mechanisms in prokaryotes, functioning by activation of effectors through the auto-phosphorylation of a conserved histidine kinase (HK) and the phosphor-transfer to its cognate partner protein, also referred to as response regulator (RR). These systems are able to sense environmental conditions and coordinate the appropriate response to ensure survival, fitness and pathogenicity (4–9).

*In silico* and functional analysis of the pneumococcal genome identified thirteen cognate HK-RR pairs and an additional orphan unpaired RR in different pneumococcal strains (10, 11). TCS in pneumococci have been associated with fitness and regulation of virulence factors, and 11 TCS are reported to contribute to pneumococcal pathogenicity (11, 12). ComDE and CiaRH, both involved in the control of competence and cell survival under stress conditions, have been studied most extensively (13–18). WalRK is another well-characterized TCS in pneumococci, featuring the only PAS (Per-Arnt-Sim) domain in *S. pneumoniae* and involved in maintenance of cell wall integrity by regulating the proteins PcsB and FabT (19–21). Furthermore, this system is the only TCS which has been shown to be essential for pneumococcal viability. However, it was proven later that this effect on viability was due to the regulation of the peptidoglycan hydrolase PcsB, whose loss-of-function leads to an unstable membrane and impaired cell viability (22, 23). Pneumococcal TCS08 (in TIGR4 genes *sp_0083* - *sp_0084*encode for RR08 and HK08) is highly homologous to the SaeRS system of *Staphylococcus aureus* (24), where it has been associated with the regulation of genes encoding α-hemolysin (*hla*), coagulase (*coa*), fibronectin (Fn) binding proteins and 20 other virulence factors (25–27). Interestingly, the SaeRS system of *S. aureus* has been shown to respond to sub-inhibitory concentrations of α-defensins and high concentrations of H_2_O_2_, suggesting a sensing mechanism responsive to host immune system molecules and membrane alterations (26, 27). In pneumococci, a previous study on TCS08 has revealed its importance for pneumococcal virulence (11). Moreover, two reports have shown a regulatory effect of the pneumococcal TCS08 on the *rlrA* pathogenicity islet (pilus-1 or PI-1) and the cellobiose phosphotransfer system (PTS) (24, 28). Hence, the initial information available about this system suggests its involvement in pneumococcal adaptation, fitness and virulence. Nevertheless, its target genes and its precise role in pneumococcal pathogenicity are yet to be defined.

## RESULTS

### Influence of TCS08 on pneumococcal growth behavior in chemically-defined medium

To investigate the effect of loss of function of TCS08 components on pneumococcal fitness, nonencapsulated *S.p*. D39 and TIGR4 parental strains and their isogenic mutants were cultured in a chemically-defined medium (CDM). All strains presented a similar growth pattern and reached similar cell densities in the stationary phase, with the exception of the TIGR4Δ*cps*Δ*rr08* mutant (Fig. 1). A steeper logarithmic phase was detected in the *rr08* mutant in TIGR4 (Fig. 1A and 1E). Additionally, the calculated growth rates of the different mutants in both D39 and TIGR4 strains suggested a significant reduction in the generation time of the *rr08* mutant in TIGR4 (Fig. 1A). The observed behavior among the TCS08 mutants in the CDM used in this study may point to strain-dependent specific effects.

**FIG 1.**
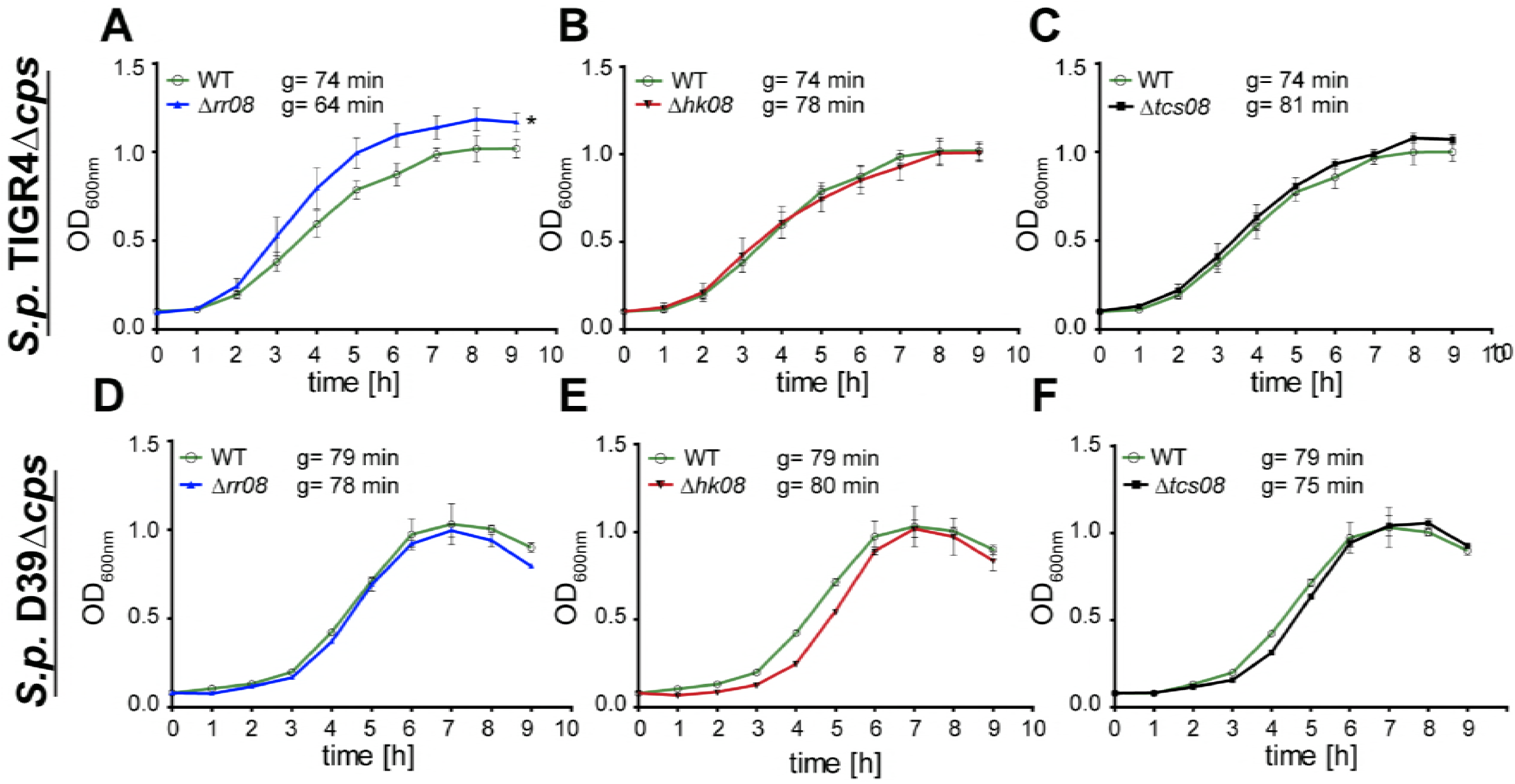
Growth behavior of pneumococcal *tcs08*-mutants. Growth in CDM of *S. pneumoniae S.p*. TIGR4Δ*cps* and *S.p*. D39Δ*cps* parental strains versus (A and D) *Δrr08-*, (B and E) *Δhk08-*and (C and F) *Δtcs08-*mutants, respectively. The symbol “g" indicates generation time. An unpaired two-tailed T-test was used with the generation times for statistics and the error bars indicate the standard deviation (SD) for n=3. The “*” symbol indicates statistical significance among the generation time of the different strains with p<0.05.

### Impact of TCS08 on TIGR4 gene expression

The initial screening for the effects of TCS08 inactivation on gene expression was conducted by microarrays using RNA samples extracted from TIGR4∆*cps* and its isogenic *rr08*-, *hk08*-, and *tcs08*-mutants grown in CDM. Genes presenting significant changes in gene expression higher than 2-fold with known functions or with functional domains were considered for further analysis. This led to the selection of 159 protein encoding genes showing significant differences in expression compared to the wild-type in at least one of the mutants deficient in RR08, HK08 or both (TCS08). Loss of the HK08 triggered the strongest changes in expression compared to the wild-type and influenced 114 genes. Differences in expression profiles of the 159 genes found in the microarray were classified by their annotated biochemical functions in 5 different categories (Fig. 2 and Table S1): (I) the largest number of genes influenced in their expression by the TCS08 was observed for genes belonging to <u>*environmental information processing (EIP)*</u>. Genes belonging to this functional class are mostly involved in membrane transport by ABC transporters and phosphotransferase systems and represented 88 genes affected by mutations in the TCS08. The strongest changes in gene expression within the EIP category were detected for the ABC transporters *aliB* (oligopeptide substrate-binding protein) and *sp_1434*, both in the *hk08* mutant. (II) The second most predominant category, with 41 genes, was the <u>*intermediary metabolism (IM)*</u>. Here, significant changes in the expression of genes involved in fatty acid (*fab*operon), carbon (cellobiose, mannitol and maltose PTS), and amino acids (*arc* operon) metabolism were seen. Indeed, the absence of the RR08 led to a significant reduction in the expression of the *arc* operon, involved in arginine uptake and utilization. In contrast, the expression of the arc-genes in the strain lacking the HK08 were upregulated. These changes observed in the expression of the *arc* operon were the most prominent within the IM category. (III) Genes reported to play a role as <u>*colonization factors (CF)*</u> accounted for 13 of the 159 genes displaying expression changes in the microarray analysis. The genes found in this group encode surface-exposed proteins involved in peptidoglycan synthesis and adhesion. Among them, the gene *sp_2136*, encoding the choline-binding protein PcpA, showed the strongest upregulation in the whole microarray analysis. The genes encoding for PavB, MucBP, PepO, PrtA and NanA displayed changes in their expression in the different *tcs08* mutants as well. These important proteins are involved in pneumococcal colonization and highlight the role of TCS08 for pneumococcal adhesion and colonization. Additionally, the lack of both components of TCS08 resulted in changes in the expression of the *rrgABC-srtC* operon, confirming the regulation of the region of diversity (RD) 4 (identified as *rlrA* or PI-1 pathogenicity islet) by TCS08. It is noteworthy, that most of these genes encode surface-displayed proteins often covalently anchored in the peptidoglycan via a transpeptidase. (IV) The fourth category encompasses genes playing a role in <u>*genetic information processing (GIP)*</u>, of which 9 genes were detected as significantly influenced by the TCS08. Genes like *rlrA*, *dnaK*, and *grpE*, are mostly involved in DNA and protein processing. Remarkably, in the absence of both components of the TCS08 a significant downregulation is seen for the positive regulator *rlrA*, involved in the expression of the PI-1. (V) The last category involves genes with an <u>*unknown function (UF)*</u>. Here, 8 genes out of the 159 identified genes presented changes in their expression in the microarray, including hypothetical lipoproteins like SP_0198 and SP_0899 (29). These proteins contain conserved lipobox motifs and are therefore also thought to be surface-exposed and might be involved in unknown fitness related processes.

**FIG 2.**
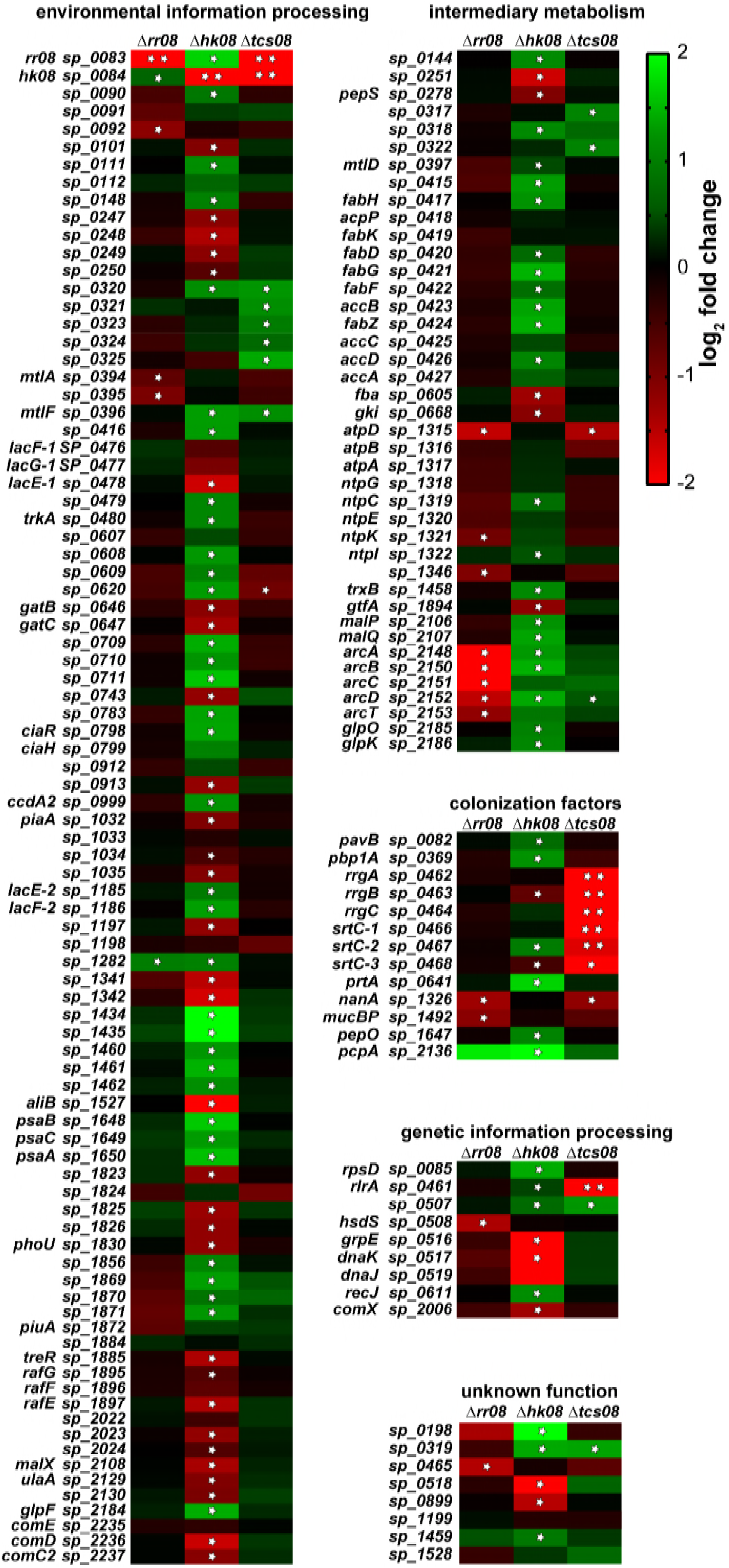
Gene expression heatmap for TIGR4 wild-type and isogenic *tcs08*-mutants. Results output for the microarray study using *S.p*. TIGR4Δ*cps* and its corresponding *tcs08*-mutants. The heat map indicates alterations in gene expression, where upregulation is indicated by green and down regulation by red colors. * indicates p-values < 0.05, ** indicates q-values < 0.05. Where q indicates False Discovery Rate statistic result.

### TCS08 is involved in the regulation of metabolic functions of *S. pneumoniae*

Results obtained by the microarray screening suggested a regulatory effect of TCS08 in the expression of genes involved in the uptake and transport of essential nutrients for *S.p*. TIGR4 such as arginine and manganese (Fig. 2 and Table S1). These metabolites/ ions are transported into the cell via specific ABC transporter systems. Of particular interest is the arginine-deiminase system (ADS), which is essential for arginine uptake and utilization in pneumococci. All genes of the *arcABCDT* operon displayed important changes in their expression in the absence of the RR or the HK08. Interestingly, these changes were not consistent in both mutants as the Δ*rr08* stain displayed a significant downregulation of this operon while the *hk08* mutant showed an upregulation (Fig. 2 and table S1). Additionally, no significant effects were observed for the *arc* operon in the Δ*tcs08* mutant. Analysis by qPCR partially confirmed the initial findings on the expression of the *arc* operon and demonstrated a strain-dependent effect for these genes. Indeed, the expression of the arginine deiminase gene *arcA* was only significantly increased in the ∆*rr08* and ∆*hk08* in TIGR4 (Fig. 3A), whereas no differences were found in D39 (Fig. 3B). Furthermore, the arginine-ornithine antiporter *arcD* (30, 31) presented a similar expression to *arcA* in TIGR4 and D39 TCS08 mutants, however the changes were not significant (Fig. 3). An additional key player in the pneumococcal fitness is the *psa* operon. This operon plays a role in the uptake of manganese and in the response to oxidative stress in the pneumococci. The analysis by microarray showed a significant increase of 2-fold in the expression of the *psa* operon for the *hk08* mutant in the TIGR4 strain (Fig. 2 and Table S1). Conversely, no statistically important effects were observed in the *psa* operon in the *rr08* and *tcs08* mutants in the same strain (Fig. 2 and Table S1). Validation of the microarray data by qPCR discovered a significant increase in the expression of *psaA* in the *rr08* mutant of D39. Surprisingly, the microarray data for the *psa* operon could not be confirmed by qPCR in TIGR4 (Fig. 3A).

**FIG 3.**
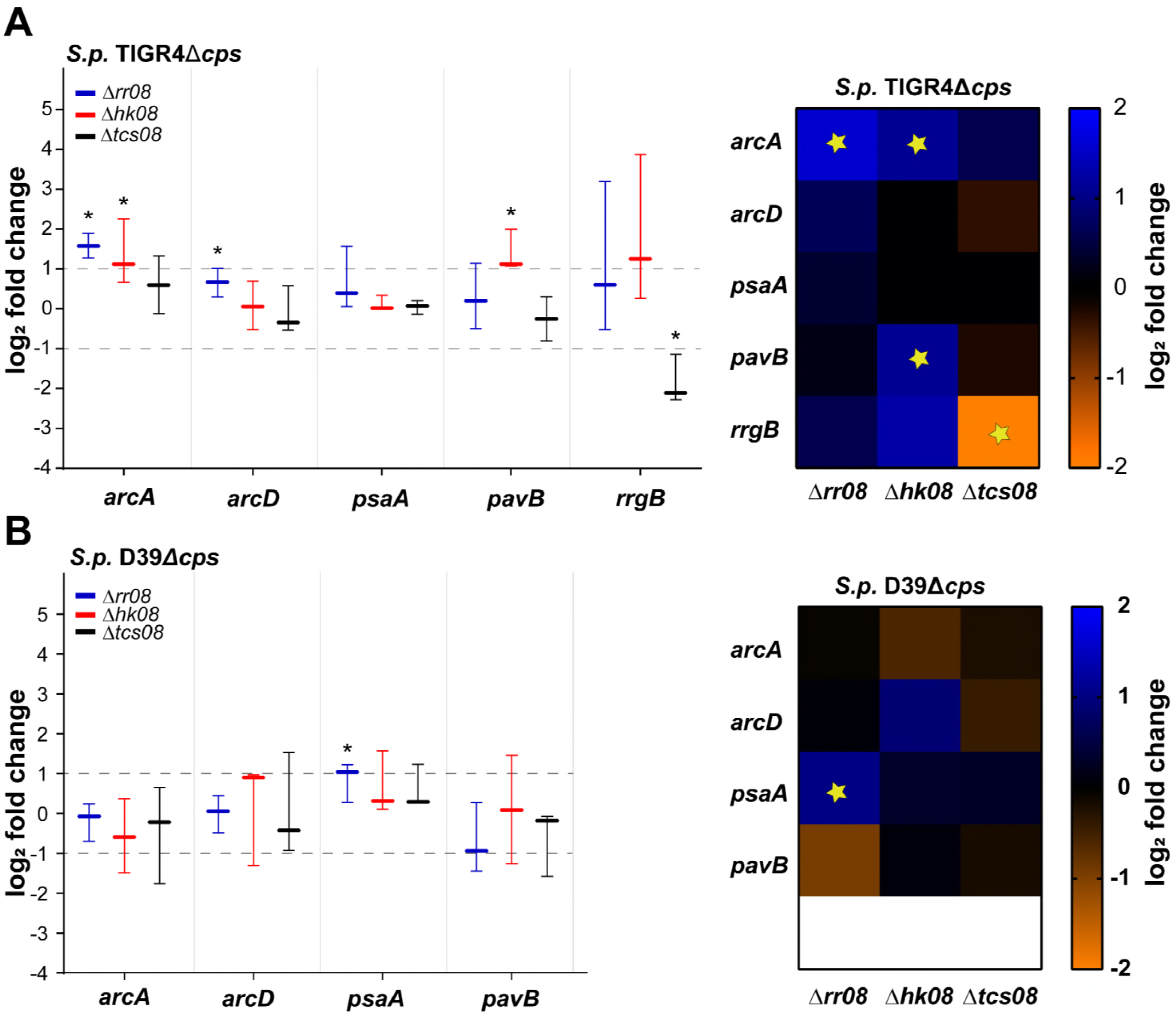
Impact of pneumococcal TCS08 on gene expression by real-time PCR. Differential gene expression in *tcs08*-mutants (–∆*rr08*, –∆*hk*08, –∆*tcs08*) analyzed by qPCR after pneumococcal cultivation in CDM. (A) *S. p*. TIGR4∆*cps* and (B) D39∆*cps*. Specific primers for the ribosomal protein S16 (*sp_0775*) were used as normalization control. Data indicates the ΔΔCt of the fold change in the graph bar and heatmap for the different *tcs08*-mutants from three independent experiments. D39Δ*cps* or TIGR4Δ*cps* wild-type were normalized to 0 and used for statistical analysis with the unpaired student’s t-test. * and * symbols indicate p-values < 0.05 in both graph and heatmap for n=3, respectively. Data are presented as boxes and whiskers with the median and 95% confidence intervals.

Immunoblot analyses of pneumococci cultured in CDM were carried out to elucidate the effect of TCS08 components on the protein levels of selected candidates from D39 and TIGR4 based on gene expression data (Fig. 4). For the ADS system, the arginine deiminase ArcA was selected as representative protein. Analysis of protein abundance of ArcA in D39 revealed a significant increase in the Δ*hk08* mutant (Fig. 4B). On the contrary, the loss of the HK08 in TIGR4 resulted in a 2-fold lower abundance of ArcA (Fig. 4A). The remaining *rr08* and *tcs08* mutants in both strains showed non-significant effects in the protein levels of ArcA. Interestingly, the results obtained for the ArcA protein in the absence of the HK08 in both strains did not reflect the transcriptome (2-fold upregulation) or qPCR results. In the case of PsaA, the immunoblot analysis confirmed a significantly higher expression of 1.5-fold in the TIGR4 *hk08* (Fig. 4A), correlating with the microarray data (Fig 2).

**FIG 4.**
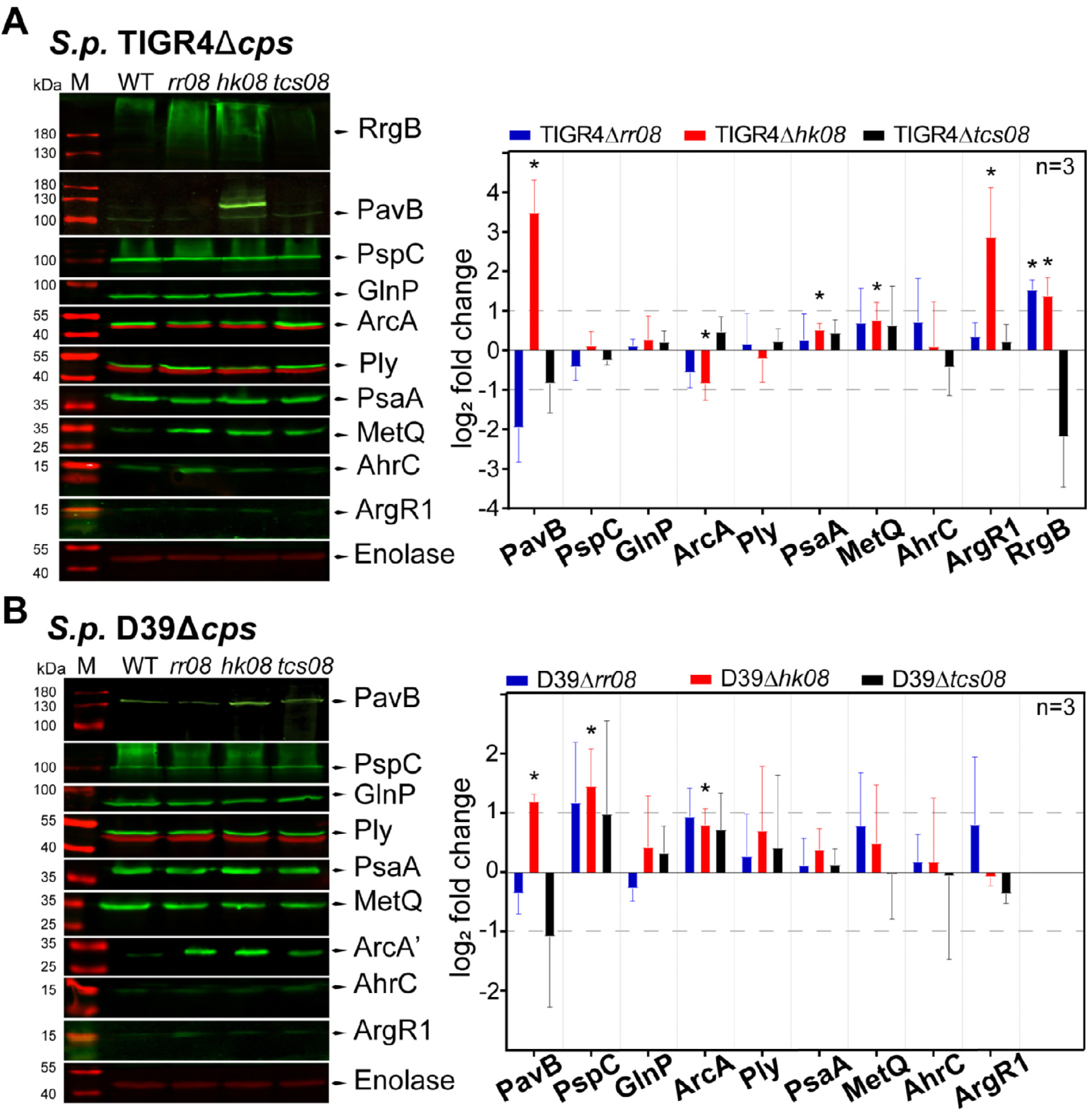
Protein expression levels in pneumococcal *tcs08*-deficient strains. Quantification of different proteins in pneumococci by immunoblotting in (A) *S.p*. TIGR4Δ*cps* and (B) *S.p*. D39Δ*cps* and their corresponding isogenic *tcs08* mutants. The unpaired student’s t-test was applied and the enolase of D39Δ*cps* or TIGR4Δ*cps* were used as reference. * indicates p-values < 0.05, n= number of biological replicates, the horizontal segmented lines indicate the 2-fold change and the error bars indicate the SD.

In a complementary approach, the surface abundance of PsaA was examined by a flow cytometric approach (Fig. 5). For D39 a non-significant increase in the surface abundance of PsaA was measured in mutants lacking both TCS08 components. The low effect of the TCS08 on PsaA observed for surface abundance correlates with the immunoblot (Fig. 4 and 5). Similarly, the increased surface abundance of PsaA in TIGR4 mutants lacking the HK08 (Fig. 5) correlated with the immunoblot and microarray analysis.

**FIG 5.**
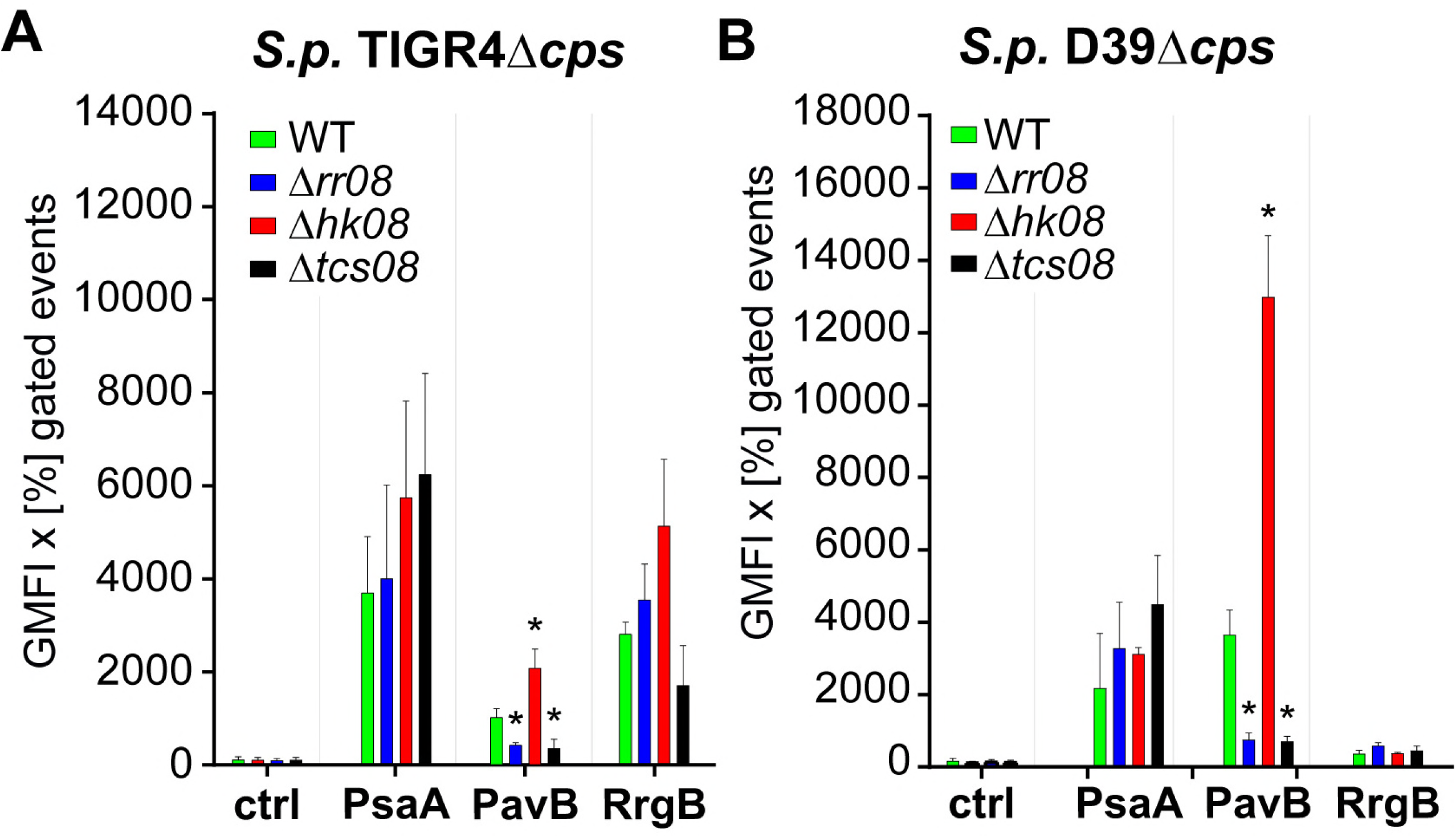
Impact of HK08 and RR08 on the abundance of pneumococcal surface proteins. The surface expression and abundance of surface proteins was analyzed by flow-cytometry in (A) *S. p*. TIGR4Δ*cps* and (B) *S. p*. D39Δ*cps* strains and their corresponding isogenic *tcs08*-mutants, all cultured in CDM. The unpaired student’s t-test was applied for the statistics and D39Δ*cps* or TIGR4Δ*cps* were used as reference accordingly. ns indicates “no significant”, * p-value<0.05 for n = 3 and the error bars indicate the SD.

### TCS08 regulates pneumococcal colonization factors

The adhesins PavB and PI-1 were shown to be regulated in the TIGR4 strain by our initial microarray analysis (Fig. 2 and Table S1) and confirmed by qPCR. Interestingly, *pavB* is a gene upstream of the 5’ region of the *tcs08* operon presenting properties of a sortase-anchored adhesin. PavB has been shown to interact with various extracellular matrix proteins and probably also directly with a cellular receptor (32, 33), thereby linking pneumococci with host cells. Similarly, the PI-1 is composed of the proteins RrgA, RrgB, and RrgC, with RrgB functioning as the backbone (34). The genes of the *pilus-1* are part of the RD4 or *rlrA* pathogenicity island and belong to the accessory genome of some pneumococcal strains and clinical isolates, including TIGR4 (35, 36). Both PI-1 and *pavB* genes presented significant changes in gene expression with an upregulation in mutants lacking the HK08 by at least 2-fold (Fig. 3). Moreover, the absence of both components of the TCS08 leads to a significantly reduced expression the *pilus-1* in TIGR4. While, no significant effect was seen for *pavB* in either the *rr08* or *tcs08* mutant in neither D39 or TIGR4 strains at the gene expression level.

On the protein level, quantifications were performed by immunoblotting (Fig. 4) and the levels of surface abundance were evaluated by flow-cytometry (Fig. 5). For PI-1, the backbone protein RrgB was used as representative. Immunoblot analysis and flow-cytometry indicated higher protein levels and surface abundance, respectively, in mutants lacking HK08 and RR08. These results are in line with gene expression analyses. Importantly, the lower protein levels of RrgB in the absence of both TCS08 components correlated with the downregulation measured by qPCR and transcriptomics (Fig. 4A and 5A). For PavB, immunoblots revealed a high impact on PavB amounts in the different mutants with a 2-fold increase in the absence of HK08 in D39 and even 10-fold in TIGR4. In contrast, the lack of either the RR08 or both components of the TCS08 procured a 2-fold decrease of PavB in both D39 and TIGR4 (Fig. 4). Similar, the surface abundance of PavB was higher in the *hk08*-mutant and lower in the *rr08*-and *tcs08*-mutants as indicated by flow cytometry (Fig. 5). Importantly, these data fit with the gene expression analysis of the mutants by microarrays.

Furthermore, an *in silico* comparison of a 300 bp upstream region of the pneumococcal gene *pavB* and the staphylococcal *saeP* and *fnbA* genes revealed the presence of a SaeR-like binding motif for *pavB* (Fig. 6). The SaeR-like binding motif is 76 bp upstream of the starting ATG of *pavB* and within its putative promotor region. In conclusion, TCS08 interferes with the regulation of adhesins and may therefore also have an impact on colonization.

**FIG 6.**
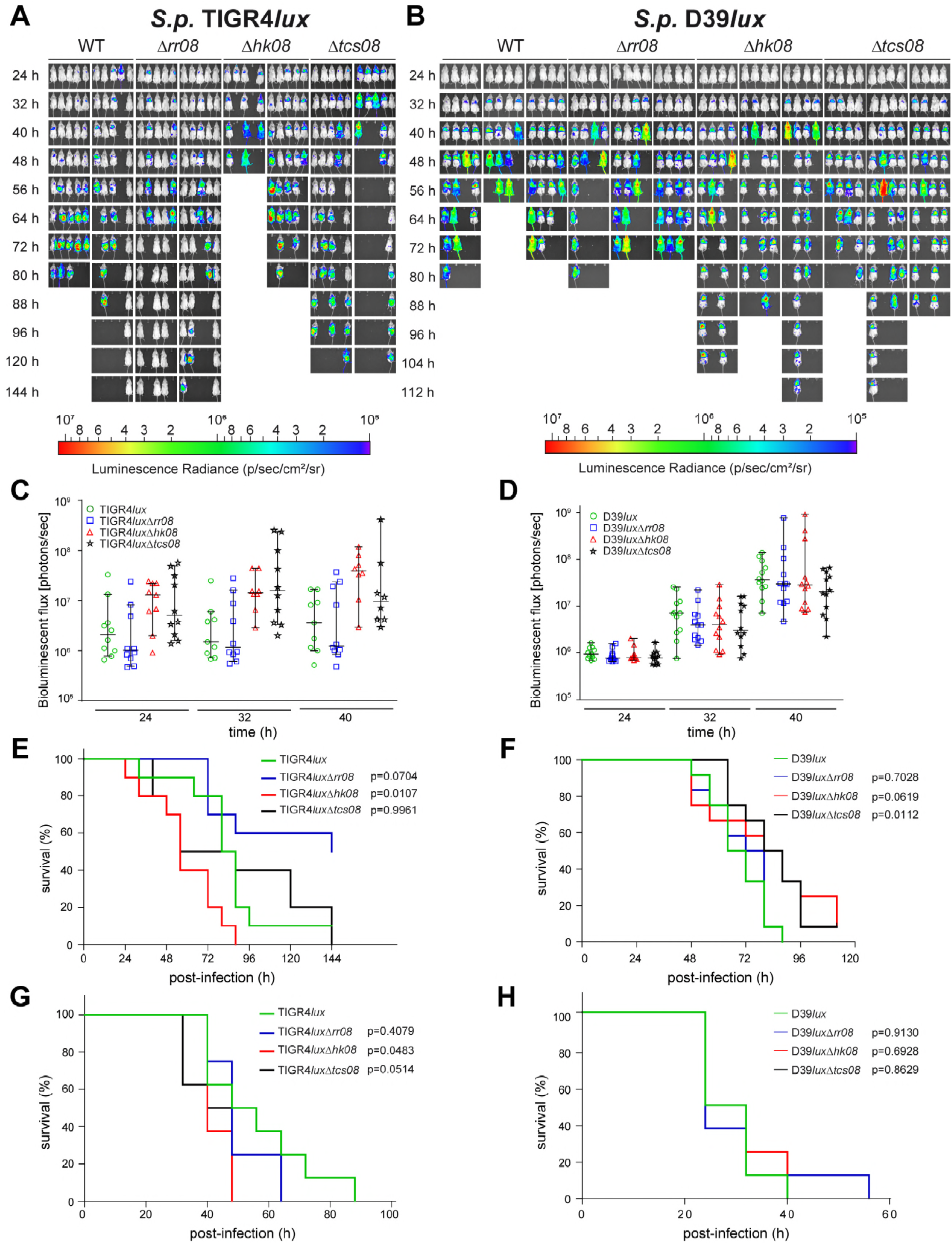
Sequence comparison of upstream regions from the genes *pavB*, *saeP* and *fnbA*. An *in silico* alignment was performed using 300 bp upstream of the pneumococcal *pavB* and the staphylococcal *saeP* and *fbnA* genes. The arrows indicate the distance upstream from the starting ATG. The bold letters in the gray boxes highlight the SaeR binding motifs in all sequences. The alignment was done using the Clustal omega tool from the EMBL-EBI. The DNA sequences were retrieved from the Kyoto Encyclopedia for Genes and Genomes (KEGG). The “*” (star) symbol indicates a conserved base pair.

### TCS08 modulation of lung infections and sepsis is strain-dependent

To assess the impact of the TCS08 or its individual components on pneumococcal colonization, lung infection or sepsis, CD-1 mice were intranasally or intraperitoneally infected with bioluminescent wild-type strains (D39 or TIGR4) and corresponding isogenic mutants. In D39, intranasal infections with mutants lacking either the HK08 or both components of the TCS08 increased the survival time of mice, thus the mutants were attenuated and represent a less virulent phenotype (Fig. 7B and F). The *rr08*-mutant of D39 showed no differences in developing lung infections (Fig. 7B and F). In the sepsis model no differences between the wild-type of D39 and its isogenic mutants were observed (Fig. 7H). Strikingly and in contrast to D39 infections, the acute pneumonia and sepsis infection models indicated a higher virulence potential of TIGR4 bacteria lacking the HK08. On the contrary, the loss of the RR08 in the TIGR4 genetic background resulted in a significantly attenuated phenotype, leading to the survival of 50% of the infected mice. No differences were observed when both components of the TCS08 were absent in TIGR4 (Fig. 7A, E and G).

**FIG 7.**
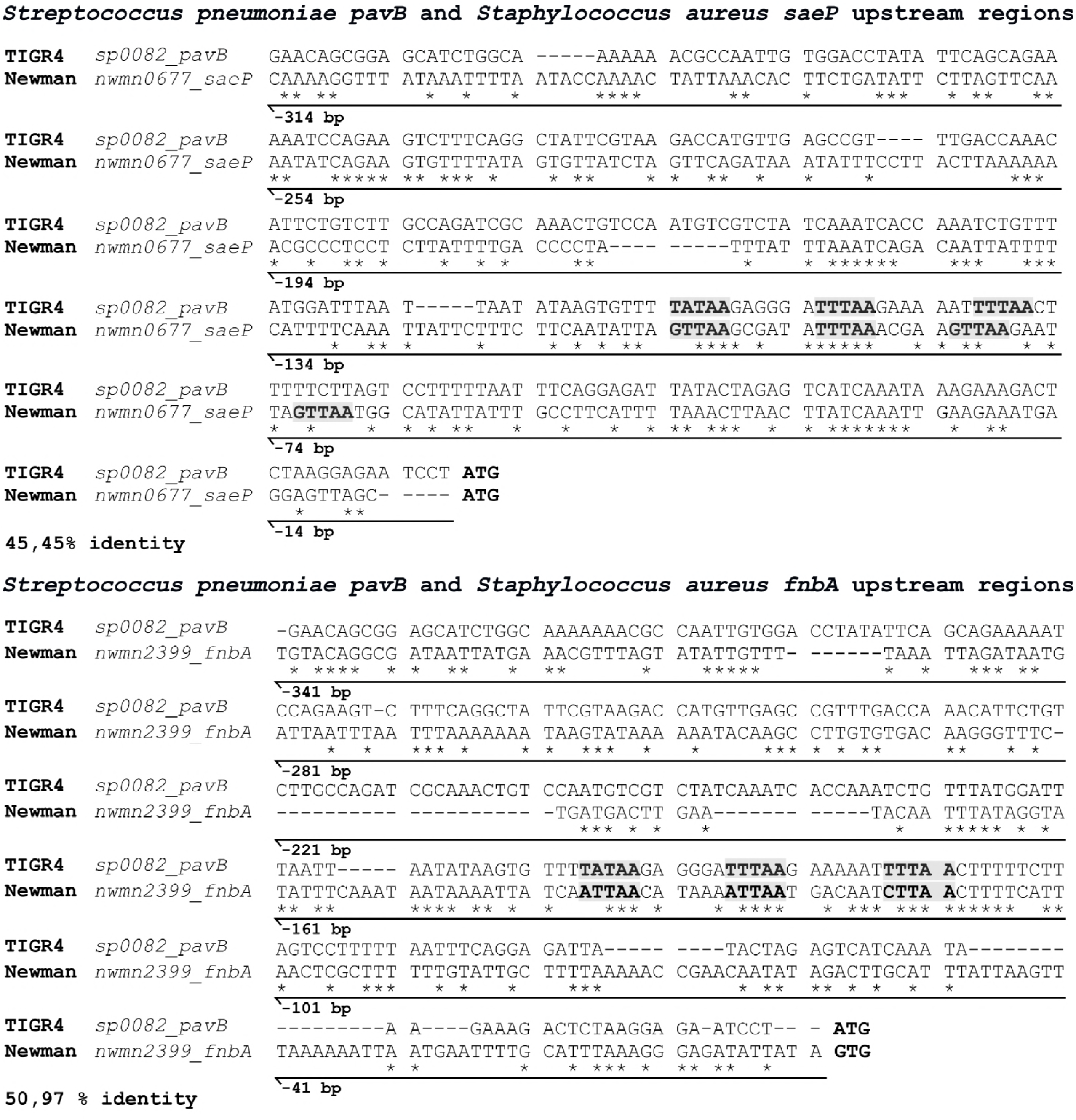
Influence of the TCS08 components on pneumococcal pathogenesis. CD-1 mice were used in the acute pneumonia model to determine the impact of the TCS08 components on virulence. Infection doses of 1×10^7^ and 7×10^7^ bacteria were applied for *S.p*. D39 and TIGR4, respectively. (A and B) Bioluminescent (*lux*) strains were used to monitor the progression of the disease *in vivo*. The results are shown as (C and D) photon flux change and (E and F) analyzed by a Kaplan-Meier plot (G and H). For the sepsis model, 1×10^3^ bacteria were used as infection dose for both wild-type strains and corresponding mutants. A log-rank test was used for the statistical test with a group size of n=12 (D39) or n=10 (TIGR4) and the error bars indicate the SD.

The impact of the TCS08 on colonization and lung infection was further investigated in the competitive mouse infection assay using the intranasal infection route. Interestingly, the wild-type TIGR4 has a lower number of recovered bacteria compared to the *rr08*-mutant, while having a significantly higher number in the nasopharynx or bronchoalveolar lavage compared to the *hk08*-mutant 24 and 48 hours post-infection (Fig. S2). Taken together, it becomes clear that the TCS08 and its individual components are essential for a balanced homeostasis, thereby maintaining pneumococcal fitness and robustness.

## DISCUSSION

The role of a subset of pneumococcal TCS in competence, physiology, and virulence has been characterized providing an initial understanding of their specific regulons (10, 12, 37). As such, TCS08 of *S. pneumoniae* has been initially identified and suggested to be important for virulence (11, 12, 37). Nevertheless, the mechanism underlying its effect on pathophysiological processes has not been elucidated before. A valid approach to estimate the regulons and effects of a TCS is to analyze the protein structures of its components. Unfortunately, only the structure of the pneumococcal RR11 and RR14 have been solved experimentally (38, 39). Nevertheless, it is possible to estimate the likely structural disposition of the remaining components by using bioinformatic tools. As such, according to the information obtained by the database SMART (Simpler Modular Architecture Research Tool, http://smart.embl-heidelberg.de/), the pneumococcal histidine kinase 08 can be classified as an intermembrane histidine kinase (IM-HK) due to its short extracellular loop. Members of this class of HK are known to respond to membrane disturbances (26). Additionally, the RR08 is classified as member of the OmpR class of response regulators, known to bind to short tandem repeats of DNA (40). Both components share a high homology and similar sequence features with the HK SaeS and RR SaeR from *S. aureus* (Fig S4) (24). Altogether, it is plausible to hypothesize that the regulatory behavior of the pneumococcal TCS08 is similar to the global virulence regulatory system SaeRS of *Staphylococcus aureus*. The staphylococcal SaeRS TCS is known to be essential for the virulence of *S. aureus* by regulating approximately 20 virulence genes such as the α-hemolysin (*hla*), fibronectin binding protein A (*fbnA*), and its own SaePQRS operon, among others (27). However, there are only a few reports regarding the control of staphylococcal fitness by the SaeRS system. One study investigated a negative regulatory effect of fatty acids on the phosphorylation of SaeS and the activation of the virulence factors controlled by SaeR (41). Our initial approach to investigate the regulatory roles of the pneumococcal TCS08 by transcriptomics discovered five main gene categories influenced by this TCS. Interestingly, we observed the most predominant regulation for genes participating in environmental information processing and intermediary metabolism (Fig. 2 and table S1). The genes grouped in these two categories are annotated as part of ABC transporters, phosphotransferase systems, and lipid biosynthesis and were found to be localized all along the pneumococcal genome (Fig. S3). The genes found to be regulated by TCS08 share an important feature, namely their localization and/or activity in the pneumococcal membrane. Additionally, several of the different PTS and ABC transporters regulated by TCS08 are involved in fitness and virulence of this pathogen. Hence, the effect of the TCS08 is more pronounced in the colonization phase of the pneumococcal life cycle. This is for example the case for the neuraminidase NanA, lipoprotein PsaA, and arginine deiminase system (ADS) (31, 42–44). Moreover, the observed regulation of the complete *fab* operon encoding enzymes for the fatty acid biosynthesis creates an important connection between the TCS08 and the sensing and responding to membrane instability (19, 45). The transporter systems affected by TCS08 are mostly essential during colonization under nutrient limiting conditions, but also in the initial stages of the diseases to take up nutrients and ensure pneumococcal fitness (Fig. 2 and 3) (46, 47).

In addition to the gene expression analysis of *tcs08*-mutants we further investigated the changes on the protein level for selected candidate proteins. Our immunoblot analyses demonstrated differences for PsaA and the arginine deiminase ArcA. Remarkably, compared to the respective wild-type strains ArcA occurred at higher protein levels in all mutants of D39 and the TIGR4 mutant lacking both the HK08 and RR08 (2-fold), while ArcA had lower protein levels in the TIGR4 mutants lacking either HK08 or RR08 (2-fold). However, only the opposite effect of deletion of *hk08* on the ArcA level was statistically significant. This is a further proof that the ADS in D39 and TIGR4 is differentially regulated as has been shown before for the stand-alone regulator ArgR2. There, the *arc* operon showed a constitutive expression in D39, while in TIGR4 gene expression was upregulated by ArgR2 (31).

It is essential that pneumococci activate their metabolic inventory when colonizing their host to ensure adaptation and fitness. As such, our results point to a role of TCS08 in the fine tuning of colonization and metabolic homeostasis as exemplified by the level of change in the expression of *pavB*, and the genes of the *pilus-1*, *fab*, and *arc* operons.

*PavB* belongs to a group of genes regulated by the TCS08 which are strongly involved in colonization by its interactions with host proteins (32, 33). These group of genes encode mostly for surface-exposed proteins associated to peptidoglycan metabolism and adherence to host cells. These genes are found grouped clockwise mostly in the first quarter of the pneumococcal genome, and transcription and replication proceed into the same direction (Fig. S3). Interestingly, the regulation of the adhesins PI-1 and PavB proteins by the pneumococcal TCS08 illustrates the high homology between the staphylococcal SaeRS and pneumococcal TCS08. Differences in gene expression of the PI-1 component genes was detected by microarray analysis (Fig. 2) and qPCR (Fig. 3) in the TIGR4 TCS08 mutants. Similarly, protein levels were also affected in the TCS08 mutants, especially in the absence of both components of the TCS08, in which a strong downregulation was detected (Fig. 4A and 5A). Our findings correlate to some extent to a previous study showing the regulation of the PI-1 by the pneumococcal TCS08 (28). For the adhesin PavB, inconsistent results were obtained for gene expression and protein abundance in the D39 strain. A minor but significant differential *pavB* gene expression was measured by microarray analysis and qPCR for TIGR4 (Fig. 2 and 3). In contrast, PavB protein levels were significantly affected in all mutants, with a 2-fold increase in the absence of the HK08 and a decrease in PavB in mutants lacking either the RR08 or both components of the TCS08 as shown by immunoblot and flow-cytometry (Fig. 4 and 5).

The staphylococcal fibronectin binding protein FbnA is weakly regulated by the SaeRS system of *S. aureus* (48), which in pneumococci correlates with the link found between TCS08 and PavB/PI-1. A direct repeat sequence (TTTAAN_7_TTTAA), similar to the imperfect SaeR binding site (GTTAAN_6_TTTAA) (49), can be found directly upstream of *pavB* (Fig. 6) suggesting that the RR08 binds directly to the *pavB* promotor region. A strong hint for the *pavB* gene regulation by the TCS08 is the higher abundance of PavB in the absence of the HK08. Surprisingly, a conserved repeat sequence TTTAAN_14_ GTTAA was found close to the *rlrA* operon and could indicate an indirect effect of the TCS08 in the regulation of the *pilus-1* via its positive regulator RlrA (Table S5). The *in silico* search for SaeR-like binding motifs among different TCS08 regulated genes indicated the presence of a variation of this binding sequence for the *cellobiose* and *arc* operons, while it was absent for the *psa* operon (Table S5). All of the genes encoded in these operons have been reported to be under the regulation of CcpA-dependent stand-alone regulators (31, 50–52). Additionally, the *psa* operon has been also shown to be under the regulation of the PsaR and TCS04 (PnpRS), which might be interplaying with the TCS08 (53, 54). This suggests either a cooperative role or a collateral effect of TCS08 and we hypothesize that the TCS08 acts as a membrane stability sensor system.

The staphylococcal SaeRS was further reported to regulate proteases and being involved in biofilm formation. Our microarray analysis showed an effect on the expression for genes encoding a putative protease domain (Fig. 2 and 3) such as the gene (*sp_0144*) possessing an Abi (abortive infective domain) with unknown function in pneumococci. Bioinformatic analysis revealed that the pneumococcal *sp_0144* is highly homologous to *spdABC* genes of *S. aureus* Newman, featuring an Abi domain. Interestingly, the SpdA, SpdB, and SpdC proteins have been reported to be involved in the deposition and surface abundance of sortase-anchored proteins in *S. aureus* (55). The gene expression of *sp_0144* (TIGR4) presented an upregulation in the hk08 mutant in TIGR4. It cannot be ruled out that the changes in SP_0144 also contribute to the protein abundance demonstrated for PavB or PI-1 when the strains lack components of the TCS08 (Fig. 3). In turn, changes in surface abundance of colonization factors will interfere with the pneumococcal virulence and /or immune evasion. However, this hypothesis was not evaluated in this study and needs experimental proof in a follow up study.

Nasopharyngeal colonization by pneumococci requires adherence to host cells and generates a foothold in the human host. Hence, the regulation of adhesins and ECM binding proteins like PavB or PI-1 represents a successful strategy of the pathogen to adapt to this host compartment. Similar, the sensing of human neutrophil peptides and membrane disruption molecules is also essential to ensure a successful colonization and immune escape phenotype. Our *in vivo* studies using pneumonia and sepsis murine models confirmed the contribution of the pneumococcal TCS08 in colonization but also virulence (Fig. 7). However, the effect is strain dependent, highlighting the role and network of different stand-alone regulators and other regulatory systems of pneumococci on the overall regulation of pneumococcal fitness and pathophysiology. Such strain-dependent effects have been also shown for additional pneumococcal TCS such as PnpRS and TCS09 (ZmpRS) (53, 56). Remarkably, a more virulent phenotype was observed for the TIGR4 mutant lacking the HK08, while the TIGR4 deficient for the RR08 displayed a decrease in virulence in the pneumonia model (Fig. 7A). In D39, the opposite effect with a slight increase in survival was observed in the absence of the HK08 in the same infection model (Fig. 7B). Additionally, the loss of function of both TCS08 components in strain D39 resulted in a significant reduction in virulence in the pneumonia model (Fig. 7B). Strikingly, this D39 attenuation was not observed in the sepsis model. Similar, the TIGR4 *rr08* mutant was also as virulent as the wild-type, despite being attenuated in the pneumonia model (Fig. 7G and 7H). In contrast, the TIGR4Δ*hk08* mutant was significantly more virulent than the wild-type in the sepsis model (Fig. 7 G). As such, our results suggest that the TCS08 is mostly involved in bacterial fitness and regulation of adhesins required for a successful colonization. Such striking difference between two representative pneumococcal strains may reflect their different genomic background and the overall versatility of pneumococci.

Interesting pathophenotypes were observed in competitive mouse infections, i.e. coinfections of the TIGR4 wild-type and its *tcs08* isogenic mutants (Fig. S2). While the pneumonia model showed an avirulent phenotype in the absence of the RR08, this mutant revealed a higher competitive index when compared to its wild-type in the coinfection assay in both, the nasopharyngeal and bronchoalveolar lavages, indicating lower numbers of the wild-type in these host compartments. In addition, TIGR4 mutants lacking either the HK08 and or both components of the TCS08 were apparently outcompeted by the wild-type (Fig. S2) despite being more virulent than the wild-type as indicated in the acute pneumonia model. A plausible explanation for this phenomenon might be that the TIGR4 mutant lacking the HK08 is rapidly progressing from the nasopharynx and lungs into the blood, and hence, low numbers are present in the nasopharynx and lavage. Similar, the absence of the RR08 impairs progressing into the blood and thus, higher numbers of the *rr08*-mutant are found in the nasopharynx. Indeed, this pneumococcal behavior post-nasopharyngeal infection can also be visualized in the bioluminescent images of the acute pneumonia model, in which the mice infected with the strain lacking the HK08 rapidly developed pneumonia and sepsis (Fig. 7A).

It is also important to mention here the mild impact of TCS08 on gene expression alterations. This suggests a role for the TCS08 as a fine tuning and signal modulation system, which is dependent on additional regulators. This hypothesis is supported by the altered gene expression of other TCS such as CiaRH and ComDE (Fig. 2 and Table S1). Such low impact on gene expression might also facilitate an explanation on the predominant role of the HK08 in controlling gene expression in pneumococci. A similar regulatory strategy has been reported for CiaRH. This system is able to control directly the expression of the protease HtrA and specific small RNAs, which in turn modulate indirectly the activity of ComDE and additional regulators (57, 58). We therefore hypothesize that the stimulus received by HK08 modulates the activity of RR08 and probably other regulators. In *Staphylococcus aureus*, the SaeRS system is also dependent on additional auxiliary proteins SaePQ (59). These proteins have been reported to interact with SaeS in order to control its phosphorylation state (59). Such systems have not yet been detected for the homologous TCS08 of the pneumococci. However, a more thorough biochemically analysis would be needed to generate a comprehensive regulatory map within pneumococcal regulators.

In conclusion, this study identified five main groups of genes influenced by the pneumococcal TCS08 in a strain-specific manner. A high number of these genes encode proteins involved in environmental signal processing, intermediary metabolism, colonization or genetic information processing. Furthermore, most of the TCS08-regulated proteins are membrane-bound and involved in nutrient transport as well as fatty acid biosynthesis. Additionally, surface-exposed PavB and PI-1 islet proteins involved in adhesion to host components were confirmed to be controlled by the TCS08. Thus, the HK08 of the TCS08 is probably sensing small molecules entering the membrane compartment of pneumococci and adapts thereby the pneumococcus to the specific environmental conditions during colonization.

## MATERIALS AND METHODS

### Bacterial strains growth conditions

*S. pneumoniae* and *E. coli* strains used in this study are listed in Table S2. Pneumococcal wild-type and isogenic *tcs08* deletion mutants were grown on Columbia blood agar plates (Oxoid) containing selection antibiotics (kanamycin 50 µg/ml and erythromycin 5 µg/ml or spectinomycin 100 µg/ml) using an incubator at 37ºC, 5% CO_2_. In liquid cultures, pneumococci were cultivated in Todd-Hewitt-broth (Roth) supplemented with 0.5% yeast extract or chemically defined medium (CDM: RPMI1640 + 2mM L-glutamine medium HyClone^TM^ GE Healthcare life sciences supplemented with 30.5 mM glucose, 0.65 mM uracil, 0.27 mM adenine, 1.1 mM glycine, 0.24 mM choline chloride,1.7 mM NaH_2_PO_4_ x H_2_O, 3.8 mM Na_2_HPO_4_, and 27 mM NaHCO_3_) using a water bath at 37°C. Recombinant *E. coli* strains were inoculated on Lysogeny Broth (LB) medium (Roth) in the presence of kanamycin (km, 50 µg/ml) at 37°C using an orbital shaker.

### Molecular techniques

The oligonucleotides and plasmid constructs used in this study are depicted in Table S3 and Table S4. The isolation of pneumococcal chromosomal DNA was achieved by using the standard phenol-chloroform extraction protocol. Briefly, *S. pneumoniae* strains were cultured in blood agar for 6 hours, transferred to new blood agar plates with antibiotics and grown for 10 hours at 37°C and 5% CO_2_. After inoculation in THY liquid medium and culture until an OD_600nm_ of 0.6 in a water bath at 37°C, the bacteria were harvested by centrifugation. The supernatant was discarded and the bacterial pellet was resuspended in TES buffer for lysis and processing. Finally, the DNA was extracted using phenol and Phenol:Chloroform:Isoamyl Alcohol (25:24:1), washed with 96% Ethanol and stored in Tris-EDTA (TE) buffer at -20°C for further use. The DNA regions needed for mutant generation and for protein production were amplified by PCR using the *pfu* proofreading polymerase (Stratagene, LaJolla, USA) and specific primers (Eurofins MWG Operon Germany) according to the manufacturer’s instructions. The annealing and extension temperatures were defined by the primers and length of the DNA inserts, respectively. The PCR products and the plasmids were purified using the Wizard^®^ SV Gel and PCR clean-up System (Promega GmbH, USA). The final constructs were confirmed by sequencing (Eurofins MWG).

### *S. pneumoniae* mutant generation

For mutant generation in D39 and TIGR4 (Δ*cps* and bioluminescent (*lux*) strains), the insertion-deletion strategy was applied by amplifying 5′ and 3′ flanking regions of *rr08*, *hk08* and the full *rr08-hk08* operon via PCR with specific primers. The genomic fragments were cloned in a pGEM-t easy vector and transformed into *E. coli* DH5α and further processed by inverse PCR using primers to delete the desired target gene and replacing it with either spectinomycin (*aad9*) or erythromycin (*erm*^*R*^) resistance gene cassettes. To achieve the deletion of the desired regions, the inverse PCR products and antibiotic cassettes were digested using specific restriction enzymes (Table S3). Finally, the deleted gene fragments encompass the following regions in each mutant: Δ*hk08* (bp 29 to 953), Δ*rr08* (bp 100 to 644) and Δ*tcs08* (bp 128 of *rr08* to bp 348 of *hk08*). Pneumococcal strains were transformed as described previously (Hammerschmidt et al., 2007 and Schulz et al., 2014) using competence-stimulating peptide (CSP) 1 (D39) or 2 (TIGR4) and cultivated in the presence of the appropriate antibiotics: kanamycin (50 µg/ml) and erythromycin (5 µg/ml) or spectinomycin (10 µg/ml). Briefly, *S. pneumoniae* strains were cultured on blood agar plates for 8 hours and a second passage was done for 10 hours in an incubator at 37ºC and 5% CO_2_. Later, the strains were inoculated in THY with an initial OD_600nm_ of 0.05 and grown in a water bath until a final OD_600nm_ of 0.1. The corresponding CSP was added and incubated at 37ºC for 15 minutes, followed by the addition of the plasmid for transformation and a heat shock treatment of 10 minutes on ice and 30 minutes at 30ºC, bacteria were allowed to grow for 2 hours at 37ºC and plated on blood agar plates with the corresponding antibiotics. The resulting *S. pneumoniae* D39 and TIGR4 *tcs08*-deficient mutants were screened by colony PCR and real-time PCR (qPCR) (Fig. S1B). Stocks were generated in THY supplemented with 20% glycerol and stored at -80°C. Individual mutants for *rr08* (*sp_0083*) and *hk08* (*sp_0084*) as well as a Δ*tcs08* (*sp_0083*+*sp_0084*) mutant were confirmed by colony PCR.

### Transcription analysis by microarrays

For the analysis of the gene expression by microarray, TIGR4Δ*cps* and its isogenic *rr*, *hk* and *tcs08* mutants were grown in CDM until an OD_600nm_ of 0.35-0.4 in triplicate. Bacterial cultures were then added to previously prepared tubes containing frozen killing buffer (20mM Tris-HCL (pH 7.5), 5mM MgCl_2_, 20mM NaN_3_) and centrifuged for 5 minutes at 10,000 g. The supernatant was completely removed and the tubes containing the pellets were immediately flash frozen in liquid nitrogen and stored at -80°C until the next step. The pellets were processed for total RNA extraction using acid phenolchloroform and DNase treatment to remove genomic DNA. The products were purified using the RNA Clean-Up and Concentration kit (NORGEN BIOTEK CORP), the quality of the RNA was determined by Agilent 2100 Bioanalyzer and the amount was quantified using a NanoDrop ND-1000 (PeqLab). 5 µg of total RNA were subjected to cDNA synthesis as described by Winter et al., (2011) (60). 100 ng of Cy3-labeled cDNA were hybridized to the microarray following Agilent’s hybridization, washing and scanning protocol (One-Color Microarray-based Gene Expression Analysis, version 6.9.1). Data were extracted and processed using the Feature Extraction software (version 11.5.1.1). Further data analysis was performed using the GeneSpring software (version 14.8). A Student´s t-test with p < 0.05, followed by a Benjamini and Hochberg false discovery rate correction with q < 0.05 were performed for the analysis.

### Gene expression analysis by qPCR

D39 and TIGR4Δ*cps* strains and their corresponding *tcs08* mutants were grown in triplicate in CDM until early-log phase and harvested for RNA isolation using the EURx GeneMatrix UNIVERSAL RNA purification kit (ROBOKLON). The RNA was checked for quality and contamination by PCR and agarose gel electrophoresis. Next, cDNA synthesis was carried out using the Superscript III reverse transcriptase (Thermofisher) and random Hexamer primers (BioRad). The obtained cDNA was checked by PCR using the same specific primers designed for the qPCR studies (Table S3). The cDNA was measured by nanodrop and stored at -20°C until further tests. For the qPCR experiments, a StepOnePlus thermocycler (Applied Biosystems) with a Syber Green master mix (BioRad) were used following the instructions for relative quantification to determine the efficiency of the primers, and as such, a reference curve was designed to be run for every gene with 5 points and concentrations ranging from 100 ng/µl to 0.01 ng/µl with 1:10 dilution steps. The StepOne software (version 2.3, Life technologies) and Microsoft^®^ Office^®^ Excel 2016 software (Microsoft) were used for the analysis. The final results are plotted as the ΔΔCT (log_2_ of the fold change of expression), with the wildtype set to 0 and compared versus its respective *tcs08* mutants. For normalization, the gene encoding the ribosomal protein S16 (*sp_0775*) was used. The results are plotted as box whiskers showing the median and 95% confidence intervals and as a heatmap.

### Protein expression by immunoblot

*S. pneumoniae* D39 and TIGR4 strains and its isogenic mutants were grown in CDM, harvested at middle-log phase and re-suspended in phosphate buffered saline buffer (PBS). A total of 2×10^8^ cells were loaded and run on a 12% SDS-PAGE and further transferred into a nitrocellulose membrane. Mouse polyclonal antibodies generated against different pneumococcal proteins and a secondary fluorescence labeled IRDye® 800CW Goat α-mouse IgG antibody (1:15000) were used to detect their expression in the WT and its isogenic mutants using the Odyssey® CLx Scanner (LI-COR). Rabbit polyclonal antibody against Enolase (1:25000) and fluorescence labeled IRDye® 680RD Goat α-rabbit IgG antibody (1:15000) were used as loading control for normalization. The quantification was performed using the Image Studio software™ (LI-COR) and the data are presented as the log_2_ of the fold change with the wild-type set to 0 and compared versus each mutant after normalization against Enolase. The Student’s t-test was used for the statistical analysis.

### Surface abundance of proteins analyzed by flow-cytometry

The expression and abundance of different surface proteins was analyzed by flow-cytometry. To detect the antigens specific primary antibodies were used in conjunction with fluorescence tagged secondary antibodies. In brief, non-encapsulated bacteria (D39Δ*cps* and TIGR4Δ*cps*) and the isogenic *tcs08*-mutants were used after growth in CDM until a final OD 0.35-0.4. Bacteria were washed with 5 ml PBS and finally resuspended in 1 ml PBS supplemented with 0,5% FCS. The bacterial cell density was adjusted to 1×10^7^ cells/ml in 1 ml of PBS/0.5% FCS/1% PFA, loaded into a 96-microtiter plate (U-bottom) and incubated for 1 hour at 4ºC. The plates were centrifuged at 3200 g for 6 minutes, the supernatant removed and bacteria were incubated for 45 minutes at 4°C with antigen specific mouse antibodies (31, 32, 61). Samples were washed twice with PBS/0.5% FCS and incubated with the goat α-mouse Alexa 488 (1/1000 dilution) antibody for 45 minutes. Thereafter, the plate was washed twice with PBS/0,5% FCS and fixed using 1% PFA in the dark at 4°C o/n. Fluorescence of the bacteria was measured using a BD FACSCalibur™ machine equipped with a log forward and log side scatter plots. The measurement of the data was conducted with the CellQuestPro Software 6.0. (BD Biosciences) collecting 50.000 events and a gated region. The results were analyzed using the freeware Flowing Software version 2.5.1 (Turku Centre for Biotechnology, University of Turku-Finland) and presented as the geometric mean fluorescence intensity (GMFI) of the analyzed bacteria population by the percentage of labeled bacteria.

### Impact of TCS08 in a murine pneumonia and sepsis models

Bioluminescent expressing *S. pneumoniae* D39*lux* or TIGR4*lux* and their isogenic mutants were grown in THY supplemented with 10% heat-inactivated fetal calf serum (FCS) until an OD_600nm_ of 0.35-0.4 and harvested via centrifugation at 3270 g for 6 min. The bacteria were resuspended in PBS and the colony forming units were adjusted for an infection dose of 1 x 10^7^ colony forming units (cfu) in 10µl or 5 x 10^3^ cfu in 200 µl per mice for the pneumonia and sepsis model, respectively. The infection process for pneumonia was carried out as follows: 8-10 weeks old 10-12 CD-1® outbread mice were arranged in groups of 5 or 4 animals per cage, respectively, and anesthetized with an intraperitoneal injection of 200 µl of Ketamin 10% (mg/ml) and 2% Rompun (dose is determined accordingly to the weight of the animals). The mice were held facing upward and 20 µl of infection dose (10 µl bacteria + 10 µl hyaluronidase (90U)) were pipetted carefully in the nostrils. Mice were allowed to inhale the drops and rest facing upwards until the anesthesia wore off. The infection dose was controlled by plating in triplicate dilutions of the bacterial solution on blood agar plates and counting the colonies. The infection was followed in real-time using the IVIS^®^ spectrum system and imaging software. Mice were controlled after the first 24 hours and every 8 hours from then on until the end of the experiment. For the sepsis model: 8-10 weeks old CD-1® outbread mice (n=8) were arranged in groups of 4 animals per cage and intraperitoneally infected with 200 µl containing 5 x 10^3^ cfu. Mice were controlled 16 hours post-infection and every 8 hours from then on until the end of the experiment. The infection dose was confirmed by plating different dilutions of the infection dose. The results were annotated using the GraphPad prism version 7.02 software and presented in a Kaplan-Meier (KM) graph. The log-rank test was used for the statistics.

Bioluminescent TIGR4 wild-type and its corresponding *tcs08*-mutants were applied in the coinfection assay. Briefly, an infection dose of 2.5 x 10^7^ cfu of wild-type and a single mutant (Δ*rr08*, Δ*hk08* or Δ*tcs08*) were mixed (1:1 ratio) and mice (n=10 CD-1) were intranasally infected. The infection dose was determined by plating serial dilutions of the infection mixture onto plates with the kanamycin or kanamycin plus erythromycin/spec to enumerate cfu of wild-type and mutant or cfu of the mutant. Mice were sacrificed after 24 and 48 hours and nasopharyngeal and bronchoalveolar lavages were performed using a tracheal cannula filled with 1ml of sterile PBS. The recovered solution was diluted and plated on blood agar plates with appropriate antibiotics (see above). Colonies were counted and recovered cfu of the wild-type and mutant determined. The competitive index (CI) was calculated as the mutant/wild-type ratio.

Values higher than 1 indicates a higher ratio of mutant bacteria, while values below 1 indicates a higher ratio of wild-type bacteria. The results were annotated using the GraphPad prism version 7.02 software and presented as scatter plots were every dot indicates 1 mouse.

## ETHIC STATEMENT

All animal experiments were conducted according to the German regulations of the Society for Laboratory Animal Science (GV-SOLAS) and the European Health Law of the Federation of Laboratory Animal Science Associations (FELASA). All experiments were approved by the Landesamt für Landwirtschaft, Lebensmittelsicherheit und Fischerei, Mecklenburg – Vorpommern (LALLFV M-V, Rostock, Germany, permit no. 7221.3-1-056/16).

## ACCESSION NUMBER

Data obtained from the microarrays analysis have been uploaded to the National Center for Biotechnology Information (NCBI) at the Gene Expression Omnibus (GEO) ArrayExpress databases at: https://www.ncbi.nlm.nih.gov/geo/ under accession number GSE108874.

## SUPPORTING INFORMATION

The supplementary information presented here includes four (5) Tables and four (4) Figures. Table S1 (results of the microarray analysis), Table S2 (list of strains and mutants), Table S3 (list of primers), Table S4 (list of plasmids) and Table S5 (*in silico* search for RR08 binding motifs). Figure S1 depicts the genomic organization and mutagenesis strategy used in this study, as well as the confirmation of the different mutants by qPCR. Figure S2 presents the competitive index obtained from the coinfection assays with the TIGR4 wildtype and its isogenic *tcs08*-mutants. Figure S3 illustrates the linear localization, orientation and category of the 159 genes obtained by the microarray study. Figure S4 illustrates the protein alignment of S. aureus SaeRS and pneumococcal TCS08.

## FUNDING INFORMATION

This work was supported by a grant from the Deutsche Forschungsgemeinschaft (DFG GRK 1870; Bacterial Respiratory Infections) in Germany, and by the Committee for Development of Research at the University of Antioquia (CODI, CIEMB-097-13) in Colombia.

## ACKNOWLEDGMENTS

We would like to acknowledge the technical work performed by Kristine Sievert-Giermann, Birgit Rietow and Gerhard Burchhardt in this study.

## AUTHORS CONTRIBUTIONS

Conceived and designed the experiments: AGM, GG, and SHA. Performed the experiments: AGM, GG, SHI, HR, UM, FV, LP, SB, NK, VK. Analyzed the data: AGM, GG, HR, UM and SHA. Writing of the manuscript: AGM, GG and SHA. Revision of the manuscript: AGM, GG, HR, UM, RB, UV and SHA.

## SUPPLEMENTS

### Figure Legends

**FIG S1** Generation of pneumococcal *tcs08*-mutants.

(A) Schematic model of the gene organization and insertion deletion mutagenesis by allelic replacement of the *tcs08* operon in *S. pneumoniae* TIGR4 as an example for all produced mutants. An *in silico* search for operon conformation identified the transcription start and terminator for the *tcs08* operon as indicated by the black arrow and lollipop, respectively. (B) Mutants were also confirmed by realtime PCR (qPCR). Specific primers were used for the *rr08* and *hk08*. Additionally, the ribosomal protein S16 (*sp_0775*) was used as control.

**FIG S2** Coinfection assay with TIGR4 wild-type and isogenic *tcs08*-mutants.

Competition assays between wild-type and TCS08 mutants were carried out in *S. p*. TIGR4. CD-1 mice were intranasally inoculated with a mixture of bioluminescent TIGR4 and each *tcs08*-mutant with an infection dose of 2.5×10^7^ of each strain. Mice were sacrificed and the samples were collected after 24 and 48 hours. Colony determination were plotted as the mutant/wild-type ratio to determine the CI for the (A) nasopharyngeal and the (B) bronchoalveolar lavages. Results are displayed as scatter plots with each dot representing one mice and the solid line indicating the median.

**FIG S3** Localization, orientation and grouping of the regulated genes by the pneumococcal TCS08.

Genes under regulation by the pneumococcal TCS08 are illustrated in a linear representation of the genome of *Streptococcus pneumoniae*. The left panel indicates the localization and orientation of each gene, where localization on the positive strand is indicated by the red color and the negative strand is indicated with the blue color. The right panel of the figure groups the genes in 5 different biochemical categories according to the characterization suggested by the databases KEGG (Kyoto Encyclopedia of Genes and Genomes) and BacMap (Bacterial Map genome atlas): green indicates Environmental information processing (EIP), yellow indicates intermediary metabolism (IM), blue indicates colonization factors (CF), pink indicates genetic information processing (GIP), and gray indicates genes of unknown function (UF).

**FIG S4** Staphylococcal SaeRS and pneumococcal TCS08 sequence alignment

Amino acid sequence alignment between the histidine kinases and response regulators of the pneumococcal TCS08 and the staphylococcal SaeRS systems. The red colored residues indicate the reported histidine and aspartate residues for SaeS and SaeR, respectively. The sequence comparison was performed using the Clustal omega tool from the EMBL-EBI and the protein sequences were retrieved using the Kyoto Encyclopedia of Genes and Genomes (KEGG). The “*” (star) symbol indicates a score of 1, the “:” (colon) indicates a score >0.5 and the single “:” (period) indicates a score >0 and <0.5.

